# Deployable CRISPR-Cas13a diagnostic tools to detect and report Ebola and Lassa virus cases in real-time

**DOI:** 10.1101/2020.05.26.116442

**Authors:** Kayla G. Barnes, Anna E. Lachenauer, Adam Nitido, Sameed Siddiqui, Robin Gross, Brett Beitzel, Katherine J. Siddle, Catherine A. Freije, Bonnie Dighero-Kemp, Samar Mehta, Amber Carter, Jessica Uwanibe, Fehintola Ajogbasile, Testimony J. Olumade, Ikponmwosa Odia, John Demby Sandi, Mambu Momoh, Hayden C. Metsky, Chloe K. Boehm, Aaron E. Lin, Molly Kemball, Daniel J. Park, Donald S. Grant, Christian T. Happi, Luis Branco, Matt Boisen, Brian M. Sullivan, Mihret Amare, Abdulwasiu Tiamiyu, Zahra Parker, Michael Iroezindu, Kayvon Modjarrad, Cameron Myhrvold, Robert F. Garry, Gustavo Palacios, Lisa E. Hensley, Stephen F. Schaffner, Andres Colubri, Pardis C. Sabeti

## Abstract

Viral hemorrhagic fevers (VHFs) remain some of the most devastating human diseases, and recent outbreaks of Ebola virus disease (EVD) ^1,2^ and Lassa fever (LF) ^3,4^ highlight the urgent need for sensitive, field-deployable tests to diagnose them ^5,6^. Here we develop CRISPR-Cas13a-based (SHERLOCK) diagnostics targeting Ebola virus (EBOV) and Lassa virus (LASV), with both fluorescent and lateral flow readouts. We demonstrate on laboratory and clinical samples the sensitivity of these assays and the capacity of the SHERLOCK platform to handle virus-specific diagnostic challenges. Our EBOV diagnostic detects both the *L* and *NP* genes, thereby eliminating the potential for false positive results caused by the rVSVΔG-ZEBOV-GP live attenuated vaccine. Our two LASV diagnostics together capture 90% of known viral diversity and demonstrate that CRISPR-RNAs (crRNAs) can be effectively multiplexed to provide greater coverage of known viral diversity. We performed safety testing to demonstrate the efficacy of our HUDSON protocol in heat-inactivating and chemically treating VHF viruses before SHERLOCK testing, eliminating the need for an extraction. We developed a user-friendly field protocol and mobile application (HandLens) to report results, facilitating SHERLOCK’s use in endemic regions. Finally, we successfully deployed our tests in Sierra Leone and Nigeria in response to recent outbreaks.

---

EBOV and LASV pose immediate, severe threats to human life and public health, as demonstrated by ongoing outbreaks of EVD in the Democratic Republic of the Congo (DRC) and LF in Nigeria. Despite their high morbidity and mortality, EVD and LF are difficult to diagnose because early symptoms, including fever, vomiting, and aches, are often indistinguishable from those of more common tropical diseases ^5–7^, and rapid point-of-care diagnostics are vital for facilitating timely clinical care and proper containment ^8^.

Despite the critical need for rapid point-of-care diagnostics for these viruses, current gold standards lack the logistical feasibility to effectively diagnose cases in endemic regions with limited infrastructure. PCR-based diagnostics are sensitive and can be rapidly developed for emerging or mutating viruses, but they are not practical as a point-of-care test as they require advanced laboratory infrastructure, a cold chain, and expensive reagents. Rapid antigen- and antibody-based tests are field deployable but are less sensitive than PCR and take longer to develop; they can also be ineffective in the early/acute stage of infection ^9–11^, a critical period for supportive care and to contain human-to-human spread.

EBOV and LASV both present distinct diagnostic challenges. The live attenuated rVSVΔG-ZEBOV-GP EBOV vaccine (Merck), currently being deployed to combat the DRC outbreak, produces EBOV GP RNA that can yield false-positive tests by GP-targeting assays, including the commonly-used GeneXpert RT-qPCR ^12–14^; similar false-positives will be a concern whenever new live attenuated vaccines are introduced for any virus. In the case of LASV, high genetic diversity in the virus through western Africa hinders development of diagnostic tools sensitive to all viral strains. The gold-standard RT-qPCR for viral detection, developed against Josiah strains derived from Sierra Leone (clade IV), has had false-positive and false-negative results when tested against recent clade II samples from Nigeria ^15,16^.

The recently developed CRISPR-based SHERLOCK (Specific High-sensitivity Enzymatic Reporter unLOCKing) platform provides a promising approach for rapidly adaptable, field-deployable diagnostics. SHERLOCK utilizes the RNA-targeting protein Cas13a for sensitive and specific detection of viral nucleic acid ^17,18^. It pairs isothermal recombinase polymerase amplification (RPA) with crRNA-guided Cas13a detection, which enables specific pairing of Cas13a with the target sequence and signal amplification via Cas13’s collateral cleavage activity ^17,19,20^. Both amplification and Cas13a-based detection are isothermal, requiring only a low-energy, single temperature heat block, compatible with point-of-care detection. SHERLOCK can be combined with HUDSON (Heating Unextracted Diagnostic Samples to Obliterate Nucleases), which inactivates pathogens and releases nucleic acid through a combined heat and chemical denaturation, eliminating the need for a column- or bead-based nucleaic acid extraction ^18^. Our recent work has shown the high sensitivity of SHERLOCK and HUDSON in detecting Zika virus and dengue virus directly from bodily fluids ^18^, allowing for a fully point-of-care diagnostic. Utilizing this system we aimed to develop a diagnostic test for EBOV and LASV that could be deployed in any setting, would require minimal processing of infectious materials, and could accurately report test results in a user friendly format.

Motivated by the increased severity and frequency of EBOV and LASV outbreaks, we describe here the development and validation of SHERLOCK assays to detect these viruses. The assays can be detected by two detection methods, either fluorescence readout or lateral flow. The more sensitive fluorescence-based system allowed us to perform extensive validation during the development of our assay, determine the length of amplification time needed for viral detection, and determine the limit of detection (LOD). The second assay, lateral flow, which we then validated further, utilizes a commercially available detection strip to provide semi-quantitative point-of-care detection of the virus.

We developed a SHERLOCK-EBOV assay to target the *L* gene of the EBOV Zaire strain, which accounts for the majority of known clinical cases of EBOV infections, including the two largest and most recent EVD epidemics ^1,2^. We used primer design applications (CATCH^21^ and ADAPT – *manuscript in prep*) to identify an optimal target within a conserved region of the *L* gene, thus avoiding potential false-positive results caused by the rVSVΔG-ZEBOV-GP EBOV vaccine (Fig 1A, Table S1 & S2). Our assay detected synthetic DNA at concentrations as low as 10 copies/μl using either fluorescent or lateral flow readout (Fig 1B-C). Because EBOV, LASV and Marburg virus (MARV) infections present with similar symptoms and have been known to co-circulate, we tested for cross-reactivity using seedstock and synthetic DNA of each virus; our assay showed no cross reactivity to either LASV or MARV (Fig 1D).

**Fig. 1.**
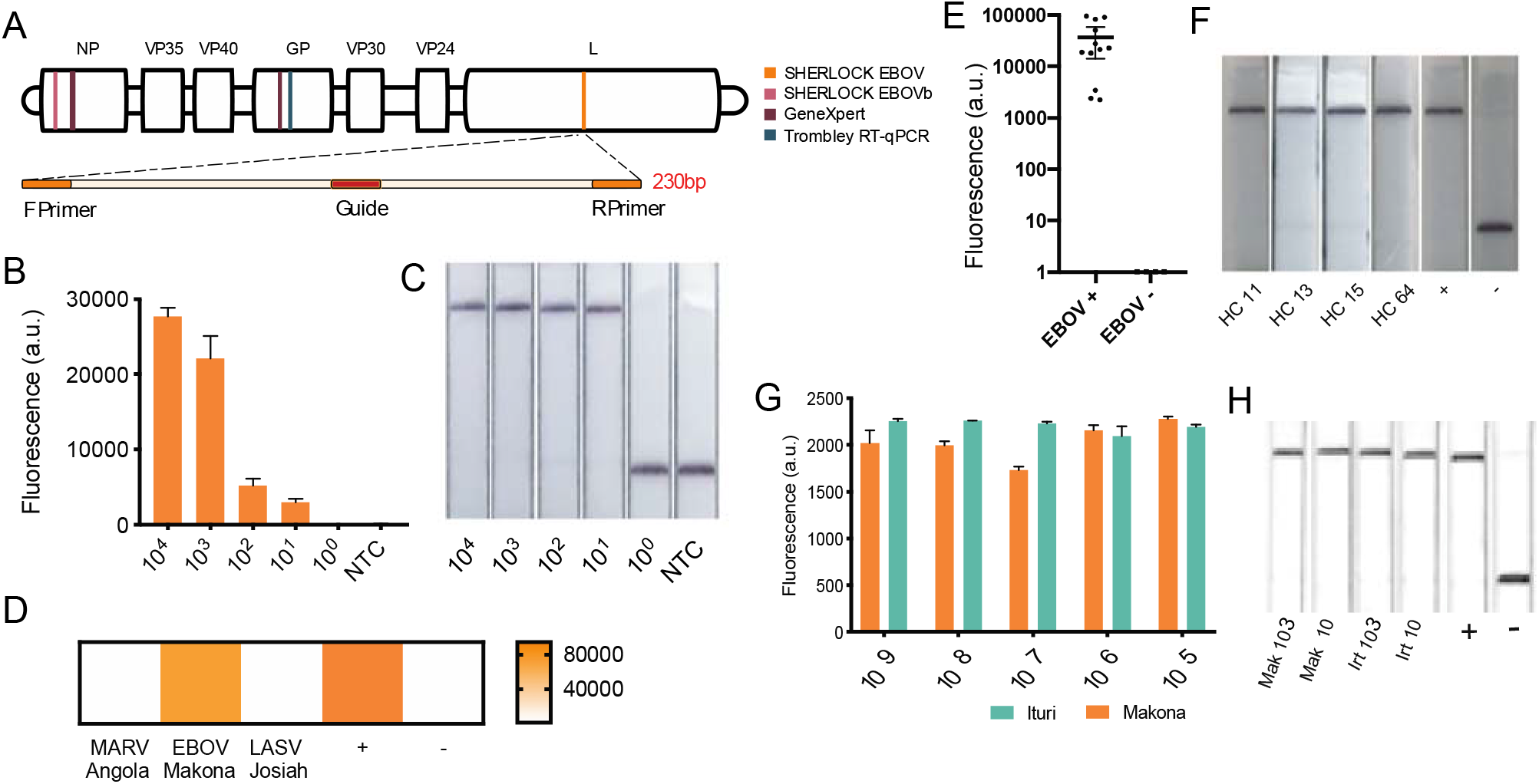
Detection of EBOV. (A) Schematic of the EBOV assay. (B) Detection of a serial dilution of EBOV synthetic DNA using (B) fluorescence and (C) lateral flow readouts for the EBOV assay. Error bars indicate 1 SD for three technical replicates. (D) Test of cross reactivity using MARV, EBOV, and LASV viral seed stock cDNA. (E) SHERLOCK testing of cDNA extracted from 12 confirmed EBOV-positive and 4 confirmed EBOV-negative samples collected from suspected EVD patients during the 2014 outbreak in Sierra Leone. Error bars indicate 95% confidence interval. (F) Four of the samples from panel E were also tested in-country using lateral flow detection. (G-H) detection of serial dilution of synthetic RNA from Ituri, DRC and Makona, Sierra Leone using (G) fluorescence and H) lateral flow readouts carried out at USAMRIID.

We validated SHERLOCK-EBOV at the Broad Institute using isolates from 16 clinical samples taken from suspected EVD patients in Sierra Leone during the 2014-2016 West Africa outbreak. For safety reasons, we tested cDNA (which is not infectious) and benchmarked the results against previously generated sequencing data ^1,22^ (Fig 1E). Of the 16 samples, 12 were positive for EBOV by sequencing, all 12 of which were positive by SHERLOCK. The 4 sequencing-negative samples were negative by SHERLOCK (100% sensitivity, 100% concordance).

Next, we deployed the assay to our collaborators in Sierra Leone to test patient samples stored from the 2014-2016 outbreak using the point-of-care lateral flow assay. This allowed us to validate and assess the practicality of the SHERLOCK assay in a setting with previously circulating EBOV and limited infrastructure. As a head-to-head comparison we identified 4 whole blood in trizol aliquots from the same patients tested in our first panel of 16 samples. These samples were stored at the Kenema Government Hospital biobank but under variable temperature conditions (−20°C with multiple power cuts). Using the same protocol as for the panel of 16 (Supp Fig 3), all four samples were positive by SHERLOCK, consistent with the results obtained at the Broad Institute (Fig 1F), despite multiple years of imperfect sample storage.

The SHERLOCK-EBOV assay was also highly efficient at detecting a more recent EBOV variant from the DRC. We tested a synthetic version of a 2018 Ebola isolate from Ituri Province (Ituri isolate 18FHV089), DRC (Fig 1G-H, Supp Fig 4A) and, as validation, a 2014 Makona isolate from Sierra Leone that underwent the same synthetic generation process (see methods) ^23,24^. Utilizing both the SHERLOCK fluorescence-based and lateral flow-based assays, the Makona and Ituri isolates were both detected at levels down to 1 copy/μl. The DRC isolate was genetically distinct from the 2014 isolate but maintained the key conserved stretch on the L gene that the SHERLOCK assay targets, which remains conserved all available genomes from the ongoing DRC outbreak.

We next developed and validated SHERLOCK assays for LASV, a challenging target because of the virus’s extreme genetic diversity, both within and especially between clades ^25^. Currently, two clades -- clade II, localized in Nigeria and clade IV, localized in Sierra Leone ^4^ -- account for over 90% of known clinical infections ^4,25^. Given this extreme genetic diversity, we designed two LASV SHERLOCK assays (Table S1 and S2). The first assay (LASV-II) targets clade II (Table S3 & S4). To ensure detection of all known genomes in this highly divergent clade, the assay contains two multiplexed crRNAs (IIA and IIB) (Fig 2A). The second assay (LASV-IV) targets clade IV; in this clade we found a more conserved region that enabled us to use a single crRNA. The LASV-II test was sensitive down to 10 copies/μl with a fluorescent readout (Fig S2A) and 100 copies/μl using lateral flow strips (Fig S2C); the LOD for LASV-IV was 100 copies/μl for the fluorescence-based assay (Fig S2B) and 1000 copies/μl for the lateral flow assay (Fig S2C). When we compared the two LASV-II crRNAs to an alternative assay with only one crRNA, the former detected LASV more quickly and identified an additional positive sample (Fig 2B). Neither the LASV-II nor the LASV-IV assay cross reacts with synthetic DNA from the other clade or with known positive patient samples from the other clade’s geographic region, nor did they cross react with EBOV or Marburg seed stock cDNA (Fig 2C and D). The LASV SHERLOCK assays are thus both species and clade specific — and therefore region specific, which can help distinguish possible imported cases from local transmission.

**Fig. 2.**
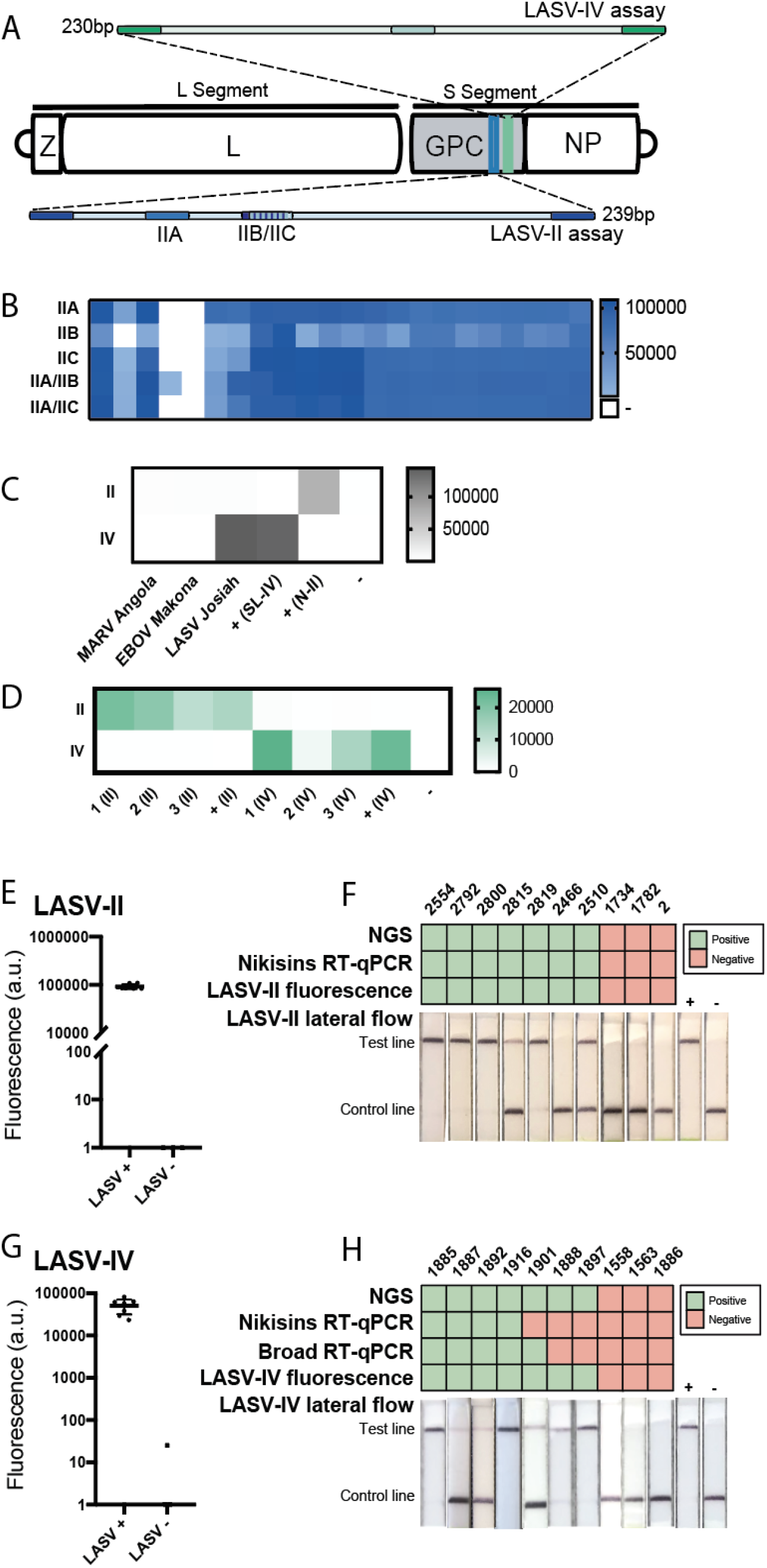
Detection of clade II and IV LASV. (A) Schematic of LASV SHERLOCK assays targeting the two most common clades of LASV (>90% known genomes): clades II (LASV-II assay) and IV (LASV-IV assay). Two primers enable target amplification via RPA, and a target-specific crRNA allows for Cas13a-based detection. For the LASV-II assay, three crRNAs were designed and tested. Two crRNAs are multiplexed to encompass the clade’s genetic diversity (IIA/IIB or IIA/IIC). (B-D) Heat maps are Fluorescence (a.u.) (B) Detection of LASV RNA from suspected LF clinical samples using crRNAs IIA, IIB, IIC, or a combination of crRNAs. (C) Test of cross reactivity between different viral species using MARV, EBOV, and LASV viral seed stock cDNA. The LASV-II and LASV-IV assays do not cross-react with MARV or EBOV seed stocks. (D) Test of cross reactivity between LASV clades using clade-specific LASV RNA from clinical samples from recent outbreaks in Nigeria and Sierra Leone. The LASV-II and LASV-IV assays provide clade-specific detection. (E) SHERLOCK testing using the LASV-II assay of RNA extracted from 7 confirmed LASV-positive and 3 confirmed LASV-negative samples collected from suspected LF patients in Nigeria during the 2018 outbreak. Lateral flow strips are shown alongside a positive control of synthetic DNA and NTC of nuclease-free water. Error bar indicates 95% confidence interval. (F) Results from panel E were compared head-to-head to those from the gold standard Nikisins RT-qPCR assay, a second Broad RT-qPCR, next-generation sequencing (NGS) (genome assembled), and lateral flow detection. (G) SHERLOCK testing using the LASV-IV assay of RNA extracted from 7 confirmed LASV-positive and 3 confirmed LASV-negative samples collected from suspected LF patients in Sierra Leone. Error bar indicates 95% confidence interval. (H) Results from panel G were compared head-to-head to those from the gold standard Nikisins RT-qPCR assay, a second Broad RT-qPCR, next-generation sequencing (NGS), and lateral flow detection.

When applied to clinical samples, the sensitivity of these LASV SHERLOCK assays was superior to that of the gold standard, RT-qPCR. We evaluated the sensitivity on a panel of 10 RNA and cDNA samples per clade, derived from suspected LF patients (Fig 2E,G). We compared SHERLOCK results using both fluorescent and lateral flow readouts head-to-head with the ‘Nikisin’ RT-qPCR assay ^26^ and benchmarked both results against sequencing data (Fig 2F,H). The LASV-II fluorescent readout was positive for all seven sequencing-positive samples, and negative for all three sequencing-negative samples (100% sensitivity, 100% concordance), as was the gold-standard Nikisin RT-qPCR. The LASV-II lateral flow readout failed to detect one sequencing-positive sample. For the LASV-IV assay, SHERLOCK performed significantly better than RT-qPCR. SHERLOCK was again positive for all seven sequencing-positive samples (100% sensitivity) while the Nikisin assay was only 40% sensitive and our in-house Broad RT-qPCR assay (Table S3), developed on all recent LASV genomes, was only 50% sensitive. The three sequencing-negative samples were negative by SHERLOCK (100% concordance). The low detection rate of clade-IV by RT-qPCRs is likely due to multiple mismatched base pairs where the primers anneal; despite this, Nikisin continues to be a primary diagnostic.

Reducing exposure to VHFs among healthcare workers is critical for the safe and effective use of diagnostic tests. For diagnostic test design, this requires ensuring that the sample is fully inactivated and, where possible, using non-invasive sample types. To this end, we combined the SHERLOCK assay with the HUDSON technique, which integrates heat inactivation with TCEP:EDTA to denature RNAses and release nucleic acid from viral particles, thus eliminating the need for RNA extraction. Furthermore, since LASV and EBOV are secreted in saliva and urine ^27,28^, HUDSON enables disease diagnosis without the need for an invasive blood draw or specialized equipment, resulting in a faster end-to-end processing time. To confirm HUDSON’s efficacy for viral inactivation, and to determine the most sensitive HUDSON protocol for SHERLOCK use, we carried out tests at the BSL4 laboratory facility at the NIH Integrated Research Facilities (IRF).

We first showed that HUDSON successfully rendered viruses inactive in three sample types. We spiked human whole blood (WB), urine, and saliva with the live EBOV Mayinga variant and confirmed that samples had viral activity using an initial plaque assay (Fig 3A). Samples underwent serial dilution to mimic variation in viral load and were then heat and chemical treated using HUDSON. We performed HUDSON using two conditions, either 95°C for 10 minutes or 70°C for 30 minutes, to determine the most effective heat-inactivation protocol. We used a standard plaque assay, two passages in Vero cells, to determine presence or absence of replication-competent virus. After HUDSON treatment, no viable virus was detected at either temperature, showing complete inactivation at all concentrations and confirming the safety of the HUDSON-SHERLOCK platform.

**Fig 3.**
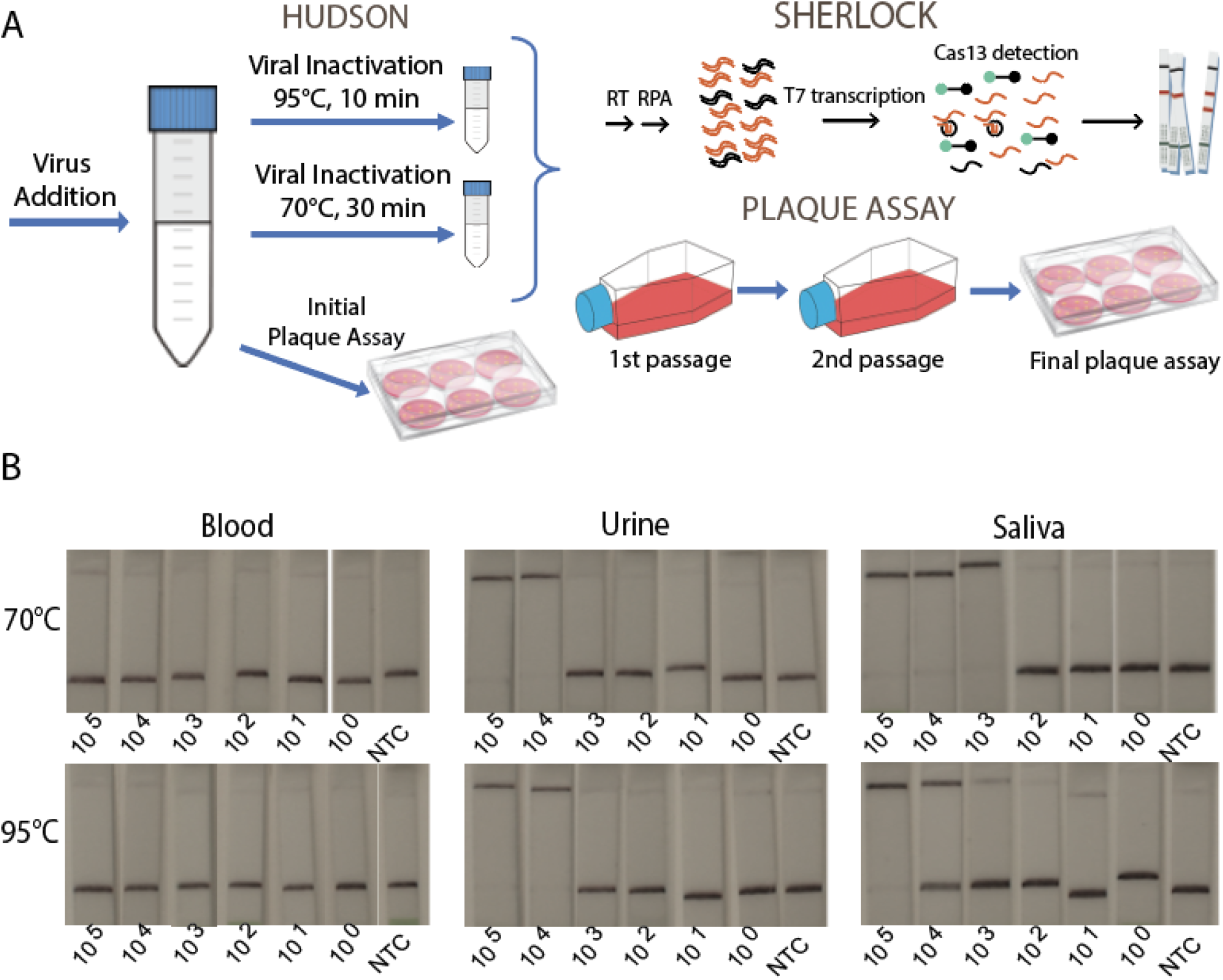
HUDSON Safety testing (A) Schematic overview of the HUDSON, SHERLOCK inactivation validation. Viral inactivation includes dilution with EDTA:TCEP and a 20min 37°C inactivation of nucleases. All final results were determined using lateral flow due to the inability to carry out appropriate fluorescent analysis in the BL4 facility. (B) Lateral flow detection of spiked blood, urine, and saliva inactivated at either 70°C or 95°C. Serial dilution shown are PFU/mL. All assays were carried out in the BL4 facility.

We then performed SHERLOCK on the HUDSON-inactivated samples to establish how HUDSON temperature conditions affect SHERLOCK’s performance. The serially diluted EBOV samples were tested by SHERLOCK using the lateral flow readout (Fig 3B), and the results were compared to the GeneXpert diagnostic (WHO, 2016). Both heat-inactivation conditions performed equally. Using our combined HUDSON-SHERLOCK method, we detected virus down to 1.1E+05 PFU/mL in WB, 1.2E+05 PFU/mL in urine and 9.4E+03 PFU/mL in saliva (Table S5). When considering cycle threshold ≤36 as definitive positives^29^ GeneXpert was more sensitive than SHERLOCK for WB (4.4E+02 PFU/mL) and urine (1.1E+02 PFU/mL), but comparable for saliva (1.1E+03 PFU/mL but for only NP detection, 9.4E+03 PFU/mL for GP detection). Ultimately, our HUDSON testing highlights the potential to safely and sensitively test saliva in suspected patients, minimizing the need for more invasive blood draws and increasing safety for healthcare workers.

The lateral flow readout of the current SHERLOCK protocol can be difficult to interpret for low concentration samples due to the correlation of band darkness with viral load. Critical for a field-deployable rapid diagnostic and surveillance tool is an easy-to-use interface that produces and reports a consistent readout free of operator bias. In addition lateral flow band strength has the potential to generate a semi-quantitative result. To exploit that potential and to facilitate accurate readout, we developed a mobile phone application ‘HandLens’, that captures and analyzes an image of one or more lateral flow strips to quantify test results and resolve ambiguous readouts (Figure 4). Using a prototype version of the HandLens app, we tested a dilution series (ranging from 10^5^-10 copies/μl) from 4 EBOV samples and compared our results to gold-standard RT-qPCR. This yielded estimates of 93% accuracy, 91% sensitivity, and 100% specificity from a total of 21 strips (Figure S5). This app can be adapted for use on any smartphone or tablet, allowing a clear, unbiased diagnostic readout.

**Fig. 4.**
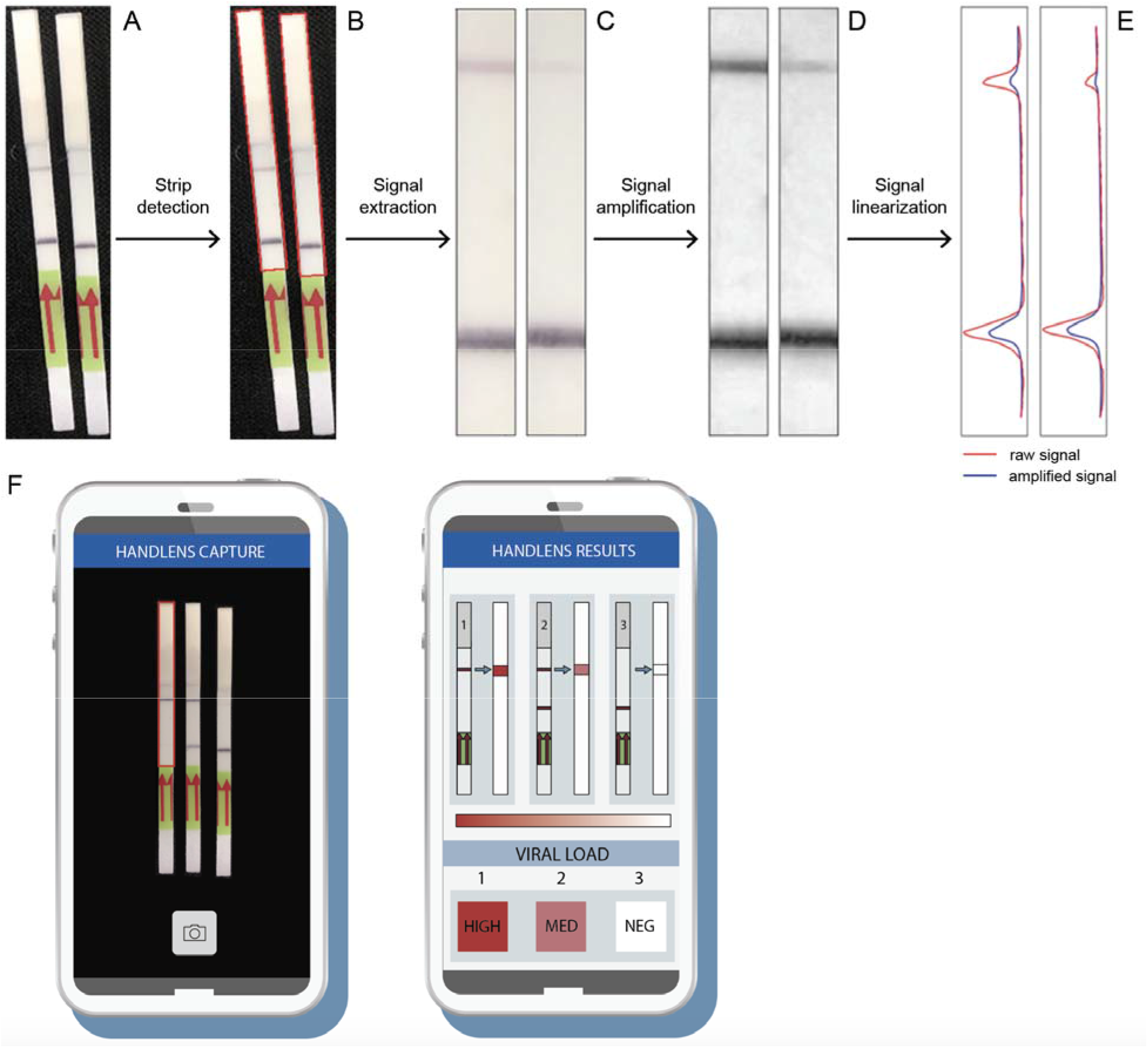
Quantification of SHERLOCK lateral flow strips using Android app prototype Internal image analysis pipeline of the SHERLOCK detector app (HandLens); (A) Images of two positive sample lateral flow strips are imported to the app (B) The relevant signal regions of the lateral flow strips are detected and demarcated by red bounding boxes. (C) Bilateral filtering is used to extract and smoothen the signal regions from the raw input image. (D) Contrast within the image are increased by applying contrast limited adaptive histogram equalization (CLAHE). (E) The signal is linearized for downstream signal processing; the red curves indicate the signal extracted after applying CLAHE, whereas the blue curves indicate the signal levels if the CLAHE step is skipped. (F) The strip reader app works by allowing the user to take a picture of the test strips where a crRNA rectangle can be used to select the control strip on the leftmost side. The raw image data is sent to a backend server that runs the signal detection algorithm and returns the binary and semi-quantitative predictions for each strip.

In summary, we have developed sensitive, specific, point-of-care CRISPR-based diagnostics for EBOV and LASV, two hemorrhagic fever viruses that pose immediate global threats. We validated these diagnostics on laboratory and patient samples, including deployment for testing in partner laboratories in Sierra Leone and Nigeria. In addition, we have shown that the HUDSON protocol not only removes the need for extraction but inactivates EBOV to allow for a safe low-tech test, and we demonstrated that non-invasive samples including saliva and urine can be used for rapid detection, eliminating the need for a blood draw and increasing safety for clinical staff testing for suspected VHF. We provide a user-friendly readout that can be documented using a mobile device to allow for greater reproducibility and immediate reporting. The HUDSON-SHERLOCK assay minimizes testing time and handling of infectious samples, reduces the cost to less than $1 USD ^30^ per sample, and can be run using only solar power or a small generator to allow for quick diagnostics in any environment, showcasing the growing capabilities of CRISPR-based diagnostics for viral detection.

## METHODS

### CLINICAL SAMPLES/ETHICS STATEMENT

All patient samples used for this study were de-identified and were obtained through studies that were evaluated and approved by the institutional review boards at the Irrua Specialist Teaching Hospital (Irrua, Nigeria), Redeemer’s University (Nigeria), Kenema Government Hospital (Sierra Leone), Sierra Leone Ministry of Health, Ministry of Health of the Democratic Republic of the Congo, and Harvard University (Cambridge, Massachusetts).

### SAMPLE PREPARATION

Patient samples were inactivated in their country of origin. Samples were then shipped to the Broad Institute or tested at the local center (Nigeria, Sierra Leone). Inactivation, RNA extraction and cDNA synthesis were carried out using the previously published methods ^31^.

### SHERLOCK ASSAY DESIGN

To design RPA primers and crRNAs, we used the computational method (ADAPT *manuscript in prep*) to identify conserved regions of the EBOV and LASV genomes. For the EBOV assay, we used an alignment based on all published sequences. The highly conserved areas of the EBOV genome allowed us to design numerous efficient crRNAs, with 2 targeting the NP-gene and 2 targeting the L-gene (Fig S1a), neither of which is expected to cross-react with the live attenuated vaccine. We developed an EBOV assay using the most efficient crRNA based on peak fluorescence and minimum time required for SHERLOCK detected to reach saturation (Fig S1b). For the LASV-IV assay, we used an alignment based on all published LASV clade IV sequences (Sierra Leone)^25^. For the LASV-II assay, we used an alignment based on all published LASV clade II sequences (Nigeria)^4,25^. We then used a 50bp sliding window to identify flanking conserved areas. We identified 21-29bp primers and a 29bp crRNA for the LASV-IV assay (Fig 1A). Due to the high diversity in clade II, one crRNA, even with up to 6 degenerate bases, did not encompass all known genomes. We identified crRNAs within the same 200bp region and tested these in tandem.

### RPA REACTIONS

All RPA reactions were carried out using the Twist-Dx RT-RPA kit according to the manufacturer’s instructions. All reactions were run for 20 minutes. Primer concentrations were 480 nM. For reactions with RNA input, Murine RNase inhibitor (NEB M3014L) was added at a concentration of 2 units/μl. For EBOV detection, the primers EBOV F and EBOV R were used. For detection of LASV clade IV, the primers LASV-IV F and R were used. For detection of LASV-II, the primers LASV-II F and R were used. For a complete list of RPA primer names and sequences, see Table S1.

### PRODUCTION OF LwaCas13a AND crRNAs

LwaCas13a was purified by Genscript. For the EBOV and LASV-IV assays, crRNAs were prepared as described ^2^. For the LASV-II assays, crRNAs were synthesized by Integrated DNA Technologies (IDT).

### CAS13a DETECTION REACTIONS

Detection reactions were performed as described ^25^. For multiplexed crRNAs in N-II assays, the total volume of crRNA was doubled. Reactions were run on a Biotek Cytation 5 multi-mode reader. All reactions were run in triplicate alongside a no-template control. Fluorescence kinetics were measured via a monochrometer with excitation at 485 nm and emission at 520 nm, with a reading every 5 minutes. EBOV assays were run for 1 hour, and LASV assays were run for 3 hours. Reactions were run at 37°C. Reported fluorescence values are specified as background-subtracted or template-specific. For EBOV detection, the crRNAs “EBOV crRNA a” and “EBOV crRNA b” were used. For detection of LASV clade IV, the crRNA “LASV-IV G1” was used. For detection of LASV-II, an equal mix of crRNAs IIA and IIB or IIA and IIC was used. For a complete list of crRNA names and sequences, see Table S1.

### LATERAL FLOW DETECTION REACTIONS

Lateral flow detection reactions were performed as described using commercially available detection strips according to manufacturer’s instructions (Milenia Hybridetect 1, TwistDx, Cambridge, UK).

### DATA ANALYSIS

For all fluorescence values, background-subtracted fluorescence was calculated by subtracting the minimum fluorescence value, which occurred between 0-20 minutes, from the final fluorescence value. For all fluorescence values reported for patient samples, found in Fig 2A-D, 3E, and 3G, target-specific fluorescence was calculated by subtracting the mean background-subtracted fluorescence of the NTC from the mean background-subtracted fluorescence of a given target with the same crRNA at the same time point.

### LOD EXPERIMENTS

To determine the sensitivity of SHERLOCK assays, assay-specific synthetically derived DNA templates were derived from clade- and virus-specific alignments ^1,4,7,22,25^. Synthetically derived DNA templates were used as input into the RPA reaction at concentrations from 10^4^ copies/μl to 1 copy/μl with a 1:10 dilution series. Each crRNA was also tested on a no input negative control. Reactions were run twice, using both the fluorescent readout and the lateral flow readout. All reactions were run in triplicate for the fluorescent readout.

### CROSS-REACTIVITY EXPERIMENTS

To assess the cross-reactivity of assays with other viruses known to cause hemorrhagic symptoms, all assays were tested on LASV (Josiah), EBOV (Makona), and MARV (Angola) viral cDNA seed stocks. Each crRNA was also tested on a positive control containing an assay-specific synthetically derived DNA template at a concentration of 10^4^ copies/μl and on no input negative control. All reactions were run in triplicate. The LASV assays were also assessed for clade-specific detection. The SL-IV assay was tested on three RNA patient samples from the LASV-II clade, and the IIA and IIB assays were tested on three RNA patient samples from the SL-IV clade. Each assay was also tested on a positive control containing an assay-specific synthetically derived DNA template at a concentration of 10^4^ copies/μl and on no input negative control. All reactions were run in triplicate.

### VALIDATION ON PATIENT SAMPLES

The LASV-IV assay was validated on a panel of 12 RNA samples collected from suspected LF patients in Sierra Leone ^4,25^. 7 of these samples were confirmed LASV-positive by antigen based RDT, ELISA IgM and RT-qPCR ^32^, 4 were LASV-negative by RDT, ELISA and RT-qPCR, and 1 did not have the full panel of tests. The assay was also tested on a positive control containing synthetically derived cDNA and a no-input control. All reactions were run in triplicate. A subset of these samples was tested using the lateral flow readout. Results were compared to RT-qPCR results and sequencing results. The LASV-II assay was validated on a panel of 12 cDNA samples collected from suspected LF patients in Nigeria during the 2018 outbreak ^4^. 9 of these samples were confirmed LASV-positive and 3 were confirmed LASV-negative. The assay was also tested on a positive control containing synthetically derived cDNA at a concentration of 10^4^ cp/μl and a no-input control. All reactions were run in triplicate. All samples were also tested using the lateral flow readout. Results were compared to RT-qPCR results and sequencing results. The EBOV assay was validated on a panel of 16 cDNA and RNA samples collected from suspected EVD patients in Sierra Leone during the 2014 outbreak ^1,22^. 12 of these samples were confirmed EBOV-positive and 4 were confirmed EBOV-negative. The assay was also tested on a positive control containing synthetically derived cDNA at a concentration of 10^4^ cp/μl and a no-input control. All reactions were run in triplicate. To validate the lateral flow readout, the EBOV assay was tested on 4 EBOV-positive cDNA samples, alongside a positive and a negative control.

### RT-qPCR EXPERIMENTS

RT-qPCR for LASV detection was performed using the Power SYBR Green RNA-to-Ct 1-step RT-qPCR kit (Thermo Fisher) according to the manufacturer’s instructions. Reactions were performed on a LightCycler 96 machine (Roche). For detection of LASV clade II, the primers Nikisins_F and Nikisins_R were used ^26^. For detection of LASV clade IV, the in-house assay including primers Broad_F and Broad_R and probe Broad_P, was also used with the TaqMan RNA-to-CT 1-Step Kit (Applied Biosystems). For a complete list of primer names and sequences, see Table S3; for a complete list of probe names and sequences, see Table S4.

### EBOV DRC EXPERIMENTS

All samples tested at USAMRIID underwent SHERLOCK as described above with the exception of the fluorescent readout which was conducted on a BioRad CFX96. The genomes of both the Makona and Ituri isolates were sequenced with Illumina technology (MiSeq for Makona and iSeq for Ituri) in-country as described in ^24^. Briefly, once the genome sequences were obtained, sub-genomic fragments were commercially synthesized and then assembled into plasmids encoding full genomes at USAMRIID. The RNAs were *in vitro* transcribed from the full genome plasmids.

### SAFETY TESTING OF HUDSON AND SHERLOCK

Viral titers for each sample were determined by plaque assay; a 6-well plate with a confluent monolayer of VeroE6 cells was infected with a predetermined volume of sample. The wells are then overlaid with a medium to ensure the monolayer health and incubated for a set amount of time. If there is live virus in the sample, this virus will infect and kill a cell, and spread cell-to-cell creating a “plaque” or a clearing of cells. “Plaques” are then visualized by a crystal violet stain and counted to determine viral titer. Samples underwent a serial dilution and then were heat-inactivated using methods described in ^18^. Briefly, a 1:100 solution of 0.5M EDTA to TCEP (Thermo) was used to decrease RNase degradation. The EDTA:TCEP was added to spiked samples at a ratio of 1 part EDTA:TCEP to 4 parts samples. First samples were heated to 37°C for 20 minutes to inactivate nucleases. The samples were then heated to 95°C for 10 minutes or 70°C for 30 minutes as described followed by the SHERLOCK assay as described above. The samples then underwent primary passaging. Samples were added to a T25 flask, incubated for 1 hour at 37°C at 5% CO_2_, then replenished with media and incubated for 7 days. 7 days post infection (dpi) pictures were taken of each flask to assess for cytopathic effect (CPE). Secondary passage was performed to assess for any residual virus particles not detected in the primary passage. All media was transferred from the T25 flask to a T75 flask incubated for 1 hour at 37°C at 5% CO_2_, replenished with media and incubated for 7 dpi and then photographed. Following the primary and secondary passaging a final plaque assay was performed to determine viral titer and to assess viral clearance or reduction. For diagnostic comparison, qPCR samples were run on the Cepheid GeneXpert platform. Sample were inactivated in Xpert lysis buffer at room temperature for 10 minutes and then run with an Xpert Ebola assay cartridge ^33^.

### HANDLENS - LATERAL FLOW READER APP

First, the signal-containing section of the lateral flow strip is detected using OpenCV’s ^34^ contour detection routines. This region is extracted and transformed into a smooth 2D image using bilateral filtering. The result is enhanced using contrast limited adaptive histogram equalization (CLAHE) ^35^, which has the property of increasing signal contrast in local regions. This signal is then linearized by integrating pixel intensity over each row. Finally, a signal is marked as positive for viral load if the signal intensity in the test band of the strip is above a certain user-defined threshold compared to the control strip, and negative if the signal intensity is too low. We also developed quantifiable signal graphs of each band and controlled for shadows and image contrast by applying a contrast-improvement algorithm.

## Acknowledgements

Opinions, interpretations, conclusions, and recommendations are those of the authors and are not necessarily endorsed by the United States Army or the National Institute of Health.

Supported by grants from the National Institute of Allergy and Infectious Diseases (NIAID), National Institutes of Health (NIH) (U19AI110818 [subaward PI: Dr. Sabeti], and the Bill and Melinda Gates Foundation (OPP1192035) (to Harvard University [Dr. Sabeti]). Dr. Barnes is supported by a NIH Fogarty K01 (K01TW010853) and Dr. Sabeti is an investigator supported by the Howard Hughes Medical Institute.

We thank the staff of the Joint West Africa Research Group, the Walter Reed Army Institute of Research, the Henry M. Jackson Foundation for the Advancement of Military Medicine, the U.S. Embassy in Abuja, Nigeria for their collaborative support and our colleagues at the Irrua Specialist Teaching Hospital, Nigeria and Kenema Government Hospital, Sierra Leone.

