## Supplementary material for "Deployable CRISPR-Cas13a diagnostic tools to detect and report Ebola and Lassa virus cases in real-time": Suppl

**Supplementary Appendix**

**Table of Contents: Supplementary Figures and Table**

Figure S1. EBOV Sherlock development, optimization and validation

Figure S2. LASV Sherlock development, optimization and validation

Figure S3. Printable field-deployable Sherlock protocol

Figure S4. DRC sample testing and cross-reactivity

Figure S5. Mobile App

Table S1. RPA primer sequences

Table S2. List of crRNA spacer sequences

Table S3. List of RT-qPCR primers

Table S4. List of RT-qPCR probes

Table S5. IRF results

**Figure S1**

(A) We developed numerous guides targeting the L (G2 and G12_13) and NP (G9 and G10_11) genes. (B) Panel shows the fluorescence of each crRNA over a 3-hour time course with a fluorescent measurement taken every 5 minutes. Input was synthetic cDNA at a concentration of 10^4^ cp/µL. Guides Ga and Gb reported the highest fluorescence compared to background. (C and D) We conducted LOD experiments on guides Ga and Gb using both fluorescent (C) and lateral flow (D) readouts. Input was synthetic cDNA at a concentration of 10^4^ cp/µL, and error bars are 1 SD based on 3 technical replicates.

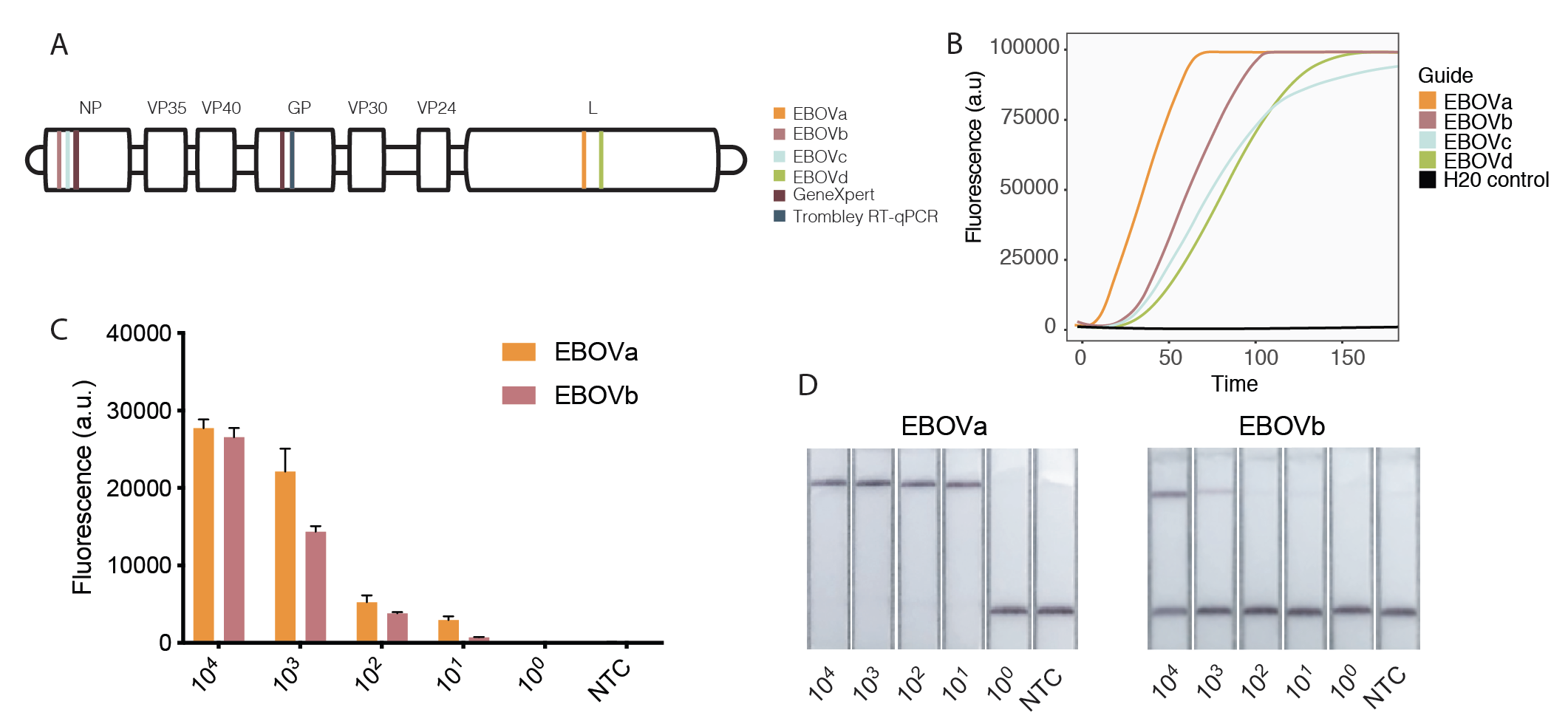

**Figure S2 LASV assay limit of detection**

Panels (A-B) show the background-corrected fluorescence of each crRNA over a 3-hour time course with a fluorescent measurement taken every 5 minutes and (C) lateral flow readout taken after a 3-hour SHERLOCK amplification. Input was synthetic cDNA at concentrations ranging from 10^5^ cp/uL-1 cp/µL, and error bars are 1 SD based on 3 technical replicates. N-IIb and N-IIa were developed to identify clade II and SL-IV was developed to identify clade IV.

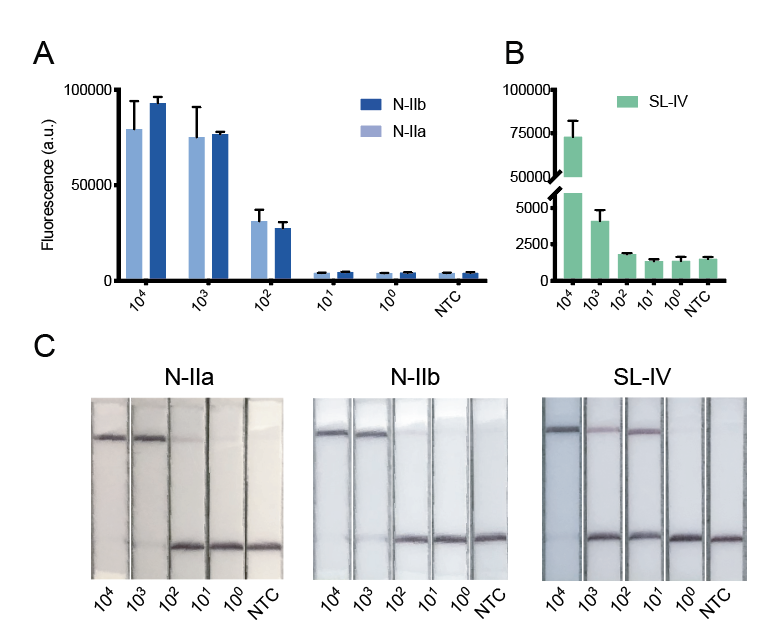

**Figure S3: Field-deployable SHERLOCK protocol**. We developed a 1-page, user-friendly SHERLOCK protocol that we shared with collaborating institutions in Sierra Leone, Nigeria, and Senegal. This field-deployable protocol simplifies SHERLOCK reactions and includes instructions for visual readout of the results.

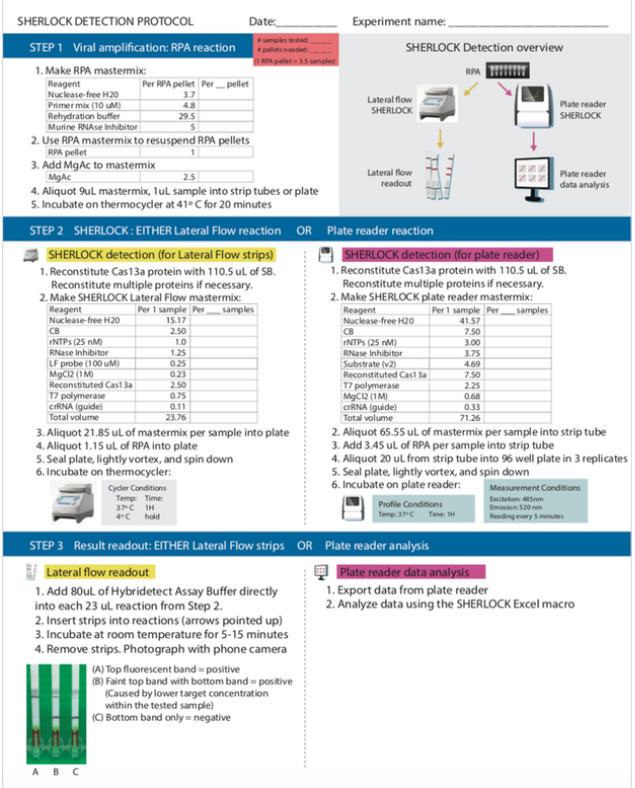

**Figure S4**

(A) Map of the DRC where red highlights the location of the Ituri outbreak. (B) Testing of other more divergent Ebola and Ebola-like isolates done at USAMRIID; There are currently six known species of Ebola: Zaire, Sudan, Tai Forest (Ivory Coast), Reston, Bombali and Bundibugyo. We were interested to determine if more divergent Ebola species would also be identified with our assay. Reston and Ivory Coast samples that were isolated at USAMRIID in 1989 and 1994, respectively, were stored in Trizol and the RNA was extracted in 2012. The Reston RNA was 30 ng/µL and the Ivory Coast RNA was 26 ng/µL but copies/µL were unknown therefore the stocks were tested using undiluted RNA (1ul) and then diluted 1:10 (0.1µL) and 1:100 (0.01µL).

A	B
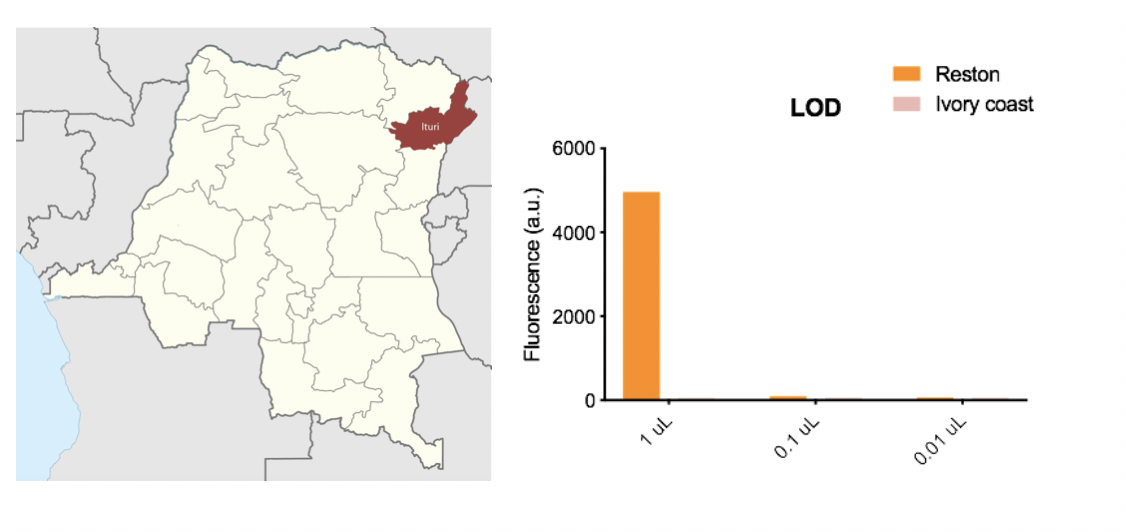

**Figure S5**

(A) Receiver Operator Characteristic Curve (ROC) generated from running the reader algorithm on from a total of 21 strips from 4 different EBOV dilution series. Of those, 17 are true positives. The AUC is 0.95, with a 95% CI of (0.88, 1.00). Using a positive classification threshold of 0.7, the algorithm exhibited the following performance indices: accuracy = 0.93, sensitivity = 0.91, and specificity = 1.00.

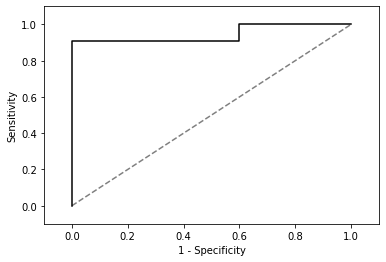

(B) The following images illustrate challenges in band detection. In the strips shown in the raw image data, the second from the right should be negative but it's classified as positive with the default classification threshold of 0.5. In CLAHE-filtered image, it is apparent that the enhancement creates a significant bump (visible in the second linearized signal plot from the top, pink line) that goes over the threshold (gray line). We currently handle these issues by increasing the classification threshold to a higher value (0.7). We plan to improve the image filtering algorithm to better handle these artifacts in the next version of the reader.

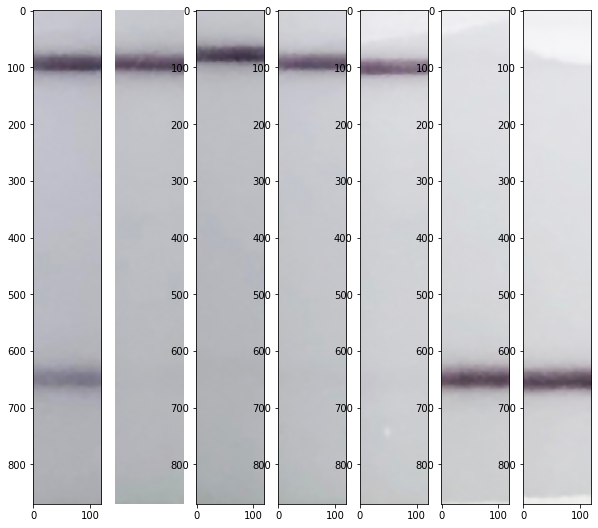

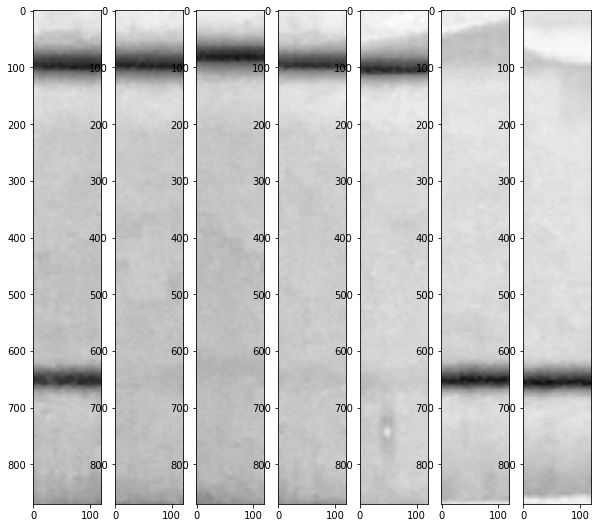

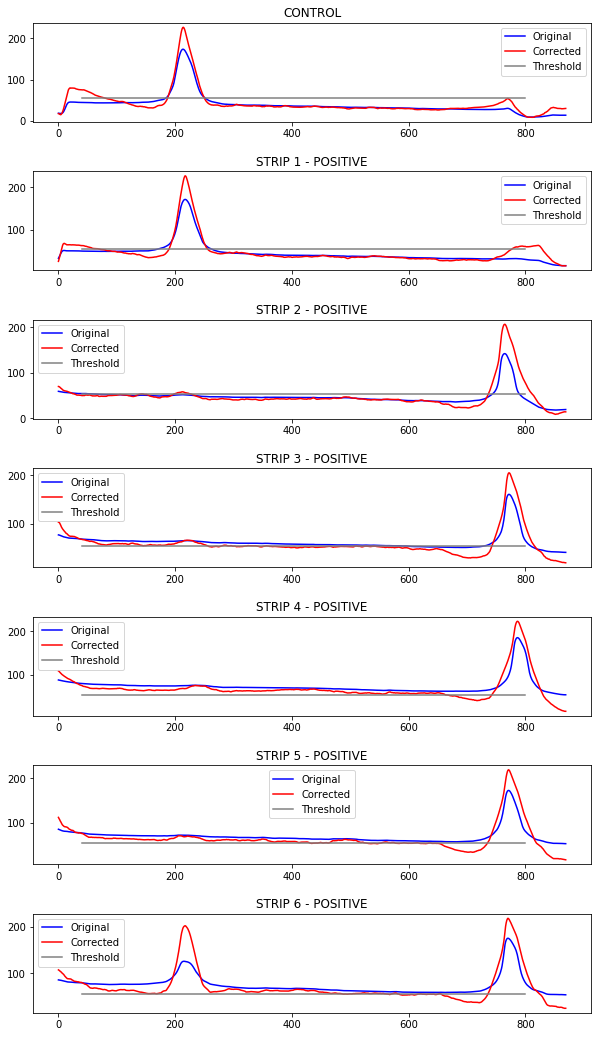

**Table S1**: List of RPA primer sequences

| RPA primer name | Sequence |
| --- | --- |
| EBOV G2 F | gaaatTAATACGACTCACTATAgggGACAGACTGAGGAARATAACATTGCAAAG |
| EBOV G2 R | CAATCATACATGGRAGTGTGGCTCCAATAA |
| EBOV G9 F | gaaatTAATACGACTCACTATAgggCAGTCAAGTAYTTGGAAGGGCACGGGTTC |
| EBOV G9 R | CTACTACCAATTTCGGAAGGAATAGACTTG |
| LASV SL-IV F | gaaatTAATACGACTCACTATAgggCATYGMATCYTTGAGRGTCAT |
| LASV SL-IV R | AGGAATCCTTATGARAACATACTCTAYAA |
| LASV N-II F | gaaatTAATACGACTCACTATAgggAAYCTYTCYGAYGCVCAYARRARGRAYCT |
| LASV N-II R | CRCCCCARGCCATYCTCATRAADGTYTG |

Table S2: List of crRNA spacer sequences

| crRNA name | Spacer sequence |
| --- | --- |
| EBOV G2 | TTTAACCCAAATAACTTGCACAGTTGAT |
| EBOV G9 | AGAACACTTGCTGCCATGCCGGAAGAGG |
| LASV SL-IV G1 | CTTCCTGTTATTGARGTYCTTGATGCAAT |
| LASV N-II G1 | CYYTRATGAGYATYATYTCAACYTTCCA |
| LASV N-II G2 | AATCARTATGARGCRATGAGYTGTGAYT |
| LASV N-II G3 | AYTTYAATCARTATGARGCRATGAGYTG |

Table S3: List of RT-qPCR primers

| Primer name | Sequence | Reference |
| --- | --- | --- |
| Nikisins_F | CCACCATYTTRTGCATRTGCCA | Nikisins *et al.* 2015 |
| Nikisins_R | GCACATGTNTCHTAYAGYATGGAYCA | Nikisins *et al.* 2015 |
| Broad_F | GATGCRGCYRAYCAYTGTG | This study |
| Broad_R | GARAACTGGCAGTGA TCTTCC | This study |

Table S4: List of RT-qPCR probes

| Probe name | Sequence | Reference |
| --- | --- | --- |
| Nikisins_P | FAM-AARTggggYCCDATgATgTgYCCWTT-BBQ | Nikisins *et al.*2015 |
| Broad_P | /56-FAM/TT YAT GAG G/ZEN/A TGG CTT GGG GTG G/3IABkFQ/ | This study |

**Table S5**

^1^Titers were determined by plaque assay. Each PFU/mL titer is accurate to ± a half-log. ^2^Not Detected – No plaques formed. ^3^Fewer than 10 plaques per well. The lower limit of Detection for the plaque assay is 100 PFU/mL ± a half-log. ^4^All GeneXpert internal controls passed.

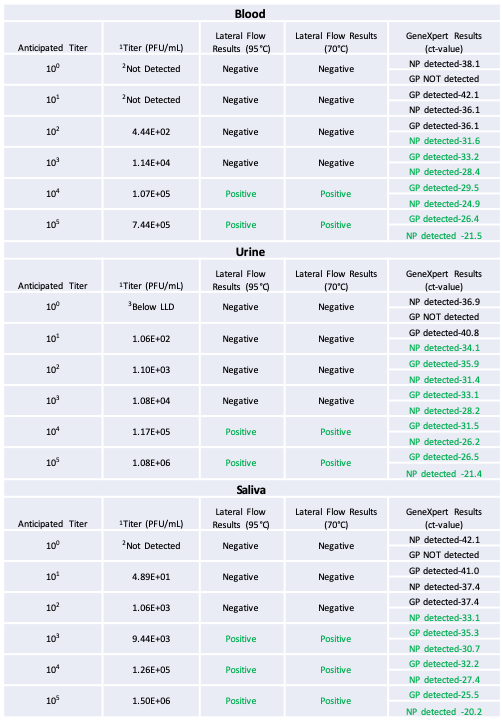
